# The capacity of the ovarian cancer tumor microenvironment to integrate inflammation signaling conveys a shorter disease-free interval

**DOI:** 10.1101/2020.04.14.041145

**Authors:** Kimberly R. Jordan, Matthew J. Sikora, Jill E. Slansky, Angela Minic, Jennifer K Richer, Marisa R. Moroney, James C. Costello, Aaron Clauset, Kian Behbakht, T. Rajendra Kumar, Benjamin G. Bitler

## Abstract

Ovarian cancer has one of the highest deaths to incidence ratios across all cancers. Initial chemotherapy is typically effective, but most patients will develop chemo-resistant disease. Mechanisms driving clinical chemo-response and -resistance in ovarian cancer are not well understood. However, achieving optimal surgical cytoreduction improves survival, and cytoreduction is improved by neoadjuvant platinum/taxane-based chemotherapy (NACT). NACT offers a window to profile pre-versus post-therapy tumor specimens, which we used to identify chemotherapy-induced changes to the tumor microenvironment. We hypothesized changes in the immune microenvironment correlate with tumor chemo-response and disease progression. We obtained matched pre- and post-NACT archival tumor tissues from patients with high-grade serous ovarian cancer (patient n=6). We measured mRNA levels of 770 genes (NanoString), and performed reverse phase protein array (RPPA) on a subset of matched tumors. We examined cytokine levels in additional pre-NACT ascites samples (n=39) by multiplex ELISA. A tissue microarray with 128 annotated ovarian tumors expanded the transcriptional, RPPA, and cytokine data by multi-spectral immunohistochemistry. In NanoString analyses, transcriptional profiles segregated based on pre- and post-NACT status. The most upregulated gene post-NACT was *IL6* (17.1-fold, adjusted p = 0.045). RPPA data were highly concordant with mRNA, consistent with elevated immune infiltration. Elevated IL-6 in pre-NACT ascites specimens correlated with a shorter time to recurrence. Integrating NanoString, RPPA, and cytokine studies identified an activated inflammatory signaling network and induced *IL6* and *IER3* (Immediate Early Response 3) post-NACT, associated with poor chemo-response and decreased time to recurrence. Taken together, multi-omic profiling of ovarian tumor samples before and after NACT provides unique insight into chemo-induced changes to the tumor and microenvironment. We integrated transcriptional, proteomic, and cytokine data and identified a novel IL-6/IER3 signaling axis through increased inflammatory signaling which may drive ovarian cancer chemo-resistance.

## INTRODUCTION

Ovarian cancer is the deadliest gynecologic malignancy in the United States, with a 63:100 death-to-incidence ratio [1]. High grade serous ovarian carcinoma (HGSOC) accounts for nearly 75% of all ovarian cancers and is the deadliest histotype, driven in large part by the high propensity of disease recurrence and rapidly acquired therapy resistance. Most patients are treated by cytoreductive surgery and combination platinum/taxane-based adjuvant chemotherapy. Though this initial chemotherapy is typically effective against residual tumor post-surgery, most patients will experience recurrence and enter a “futile cycle” of response and recurrence with additional courses of chemotherapy. This ultimately leads to the development of wholly chemo-resistant disease. Our understanding of clinical chemo-resistance in ovarian cancer is limited, as access to treated tumor tissue after the initial cytoreductive surgery and subsequent chemotherapy is uncommon. This lack of access to treated tumor hinders our ability to define how ovarian tumors respond to and adapt to therapy, and to identify predictors and drivers of therapy resistance and recurrence.

Though our mechanistic understanding of chemo-response is limited, successful surgical removal of tumor remains one of the best predictors of overall survival. Based on this, patients with HGSOC may undergo a diagnostic laparoscopy to determine the extent of disease burden and likelihood for successful optimal cytoreduction (i.e. Fagotti Scoring [2, 3]). Extensive, unresectable disease is defined by a Fagotti score of greater than or equal to 8 (e.g. due to involvement of the small bowel mesentery), and is strongly associated with decreased time to recurrence and poor patient outcomes. However, tumor resectability can be improved by the use of <u>n</u>eo<u>a</u>djuvant <u>c</u>hemo<u>t</u>herapy (NACT) with interval cytoreductive resection. NACT consists of three cycles of carboplatin/paclitaxel, then an interval surgical debulking of the remaining disease, followed by another three carboplatin/paclitaxel cycles. NACT improves rates of optimal surgical cytoreduction in patients with advanced disease,[4–6], therefore patients with advanced HGSOC are likely to receive NACT. Importantly, the carboplatin/paclitaxel chemotherapies used in NACT also remain the standard upfront care for adjuvant therapy. Since the NACT paradigm provides access to the tumor site before and after chemotherapy, NACT for HGSOC presents a unique opportunity to investigate the biological mechanisms of tumor response to current standard of care adjuvant chemotherapy.

Though studies of HGSOC tumor response to NACT are emerging, studies to date have identified a common thread in a key role for immune/inflammatory signaling in chemo-response and –resistance. Böhm and colleagues examined matched pre and post-treated HGSOC tumors to define chemotherapy-induced changes in the immune system and characterized expression of immune checkpoint receptors that attenuate immune responses [7]. They reported that following NACT, an increase in tumor infiltration of immunosuppressive T regulatory cells and elevated T cell expression of checkpoint receptors (PD1 and CTLA4) correlate to a poorer prognosis [7]. Lo and colleagues reported a similar finding: while chemotherapy promotes an immune response in most cases, this immune response does not circumvent an established immunosuppressive microenvironment at baseline [8]. Beyond NACT, several studies reported that the cytokine IL-6 significantly contributes to the immunosuppressive microenvironment in HGSOC, and is associated with disease progression and poor response to chemotherapy [9–13]. These observations suggest a key role for immune cytokines in modulating tumor response to chemotherapy, but the efficacy of immunotherapies in patients with HGSOC remains limited. Notably, the only approved immunotherapy for treatment of ovarian cancer is an anti-PD-1 directed therapy (Pembrolizumab [14, 15]). Greater mechanistic understanding of the role of the tumor immune microenvironment may allow us to overcome chemo-resistance in HGSOC and improve patient outcomes.

We hypothesized that integrating multiple profiling methods of both the tumor and tumor microenvironment pre- and post-NACT would identify predictors and drivers of chemotherapy response. To address this, we identified a series of matched archival tumor samples pre- and post-NACT from 6 patients with HGSOC. Using these samples, we defined the transcriptomic effects of NACT using the Nanostring PanCancer Immuno-Oncology 360 panel, a series of 770 genes associated with the tumor immune microenvironment. NACT-driven transcriptomic changes were linked to upstream signaling using reverse phase protein array (RPPA) analyses of a subset of samples. Additionally, we examined the cytokine milieu in pre-treatment ascites samples from 39 patients with HGSOC. Integrating these multi-omic analyses via network analysis and comparing data against different platforms (e.g. RNA expression and immunohistochemistry) identified an immune microenvironment signaling pathway associated with decreased time to recurrence, including a cytokine and associated receptor tyrosine kinase, downstream transcription factor targets, and key target genes. These data suggest that the magnitude of chemotherapy-induced remodeling of the tumor microenvironment provides a unique predictor of disease recurrence, and supports that multi-omic profiling of pre- and post-NACT samples is a powerful model for understanding chemotherapy response and resistance in HGSOC.

## MATERIALS AND METHODS

### Gynecologic Tumor and Fluid Bank (GTFB)

Patients in this study were consented under the GTFB at the University of Colorado Anschutz Medical Campus (COMIRB#07-935). Eligible patients included those being treated for gynecologic malignancy, particularly HGSOC. Patient specimens were de-identified and uniformly processed into formalin-fixed paraffin embedded tissue blocks. A portion of the tumors was also flash-frozen and stored at −80°C. Ascites was centrifuged at 200 x g for 7 minutes and the fluid and cellular components were cryopreserved.

### Multiplex enzyme-linked immunosorbent assay

The concentration of cytokines and chemokines was assessed in cryopreserved ascites fluid using the V-PLEX Pro-inflammatory Panel 1 kit according to the manufacturer’s instructions (Meso Scale Discovery). Briefly, ascites fluid was diluted 2-fold in Diluent 2, incubated for 2 h while shaking at 600 rpm at RT, then incubated for 2 h with detection antibody prepared in Diluent 3 while shaking at 600 rpm at RT. The plates were washed in between steps using an automated plate washer (Biotek Instruments). Electrochemiluminescence values were read in 2x Read Buffer on the Meso QuickPlex SQ 120 (Meso Scale Discovery). Sample concentrations were extrapolated by comparison to the provided standard curve of diluted calibrators.

### RNA Extraction

Formalin-fixed paraffin-embedded (FFPE) tissue blocks were acquired through the Department of Pathology (COMIRB#18-0119). Tissue blocks were examined by a board-certified pathologist to confirm the presence of tumor tissue. FFPE tissue blocks were sectioned, and 10 micron tissue-containing paraffin scrolls were collected. RNA was extracted using the High Pure FFPET RNA Isolation kit (Roche) according to the manufacturer’s instructions and eluted in 40 μl Elution Buffer. RNA quantity and quality was assessed using a RNA Screentape on a TapeStation 4150 (Agilent). The concentration of RNA was determined by comparison to the RNA ladder (average 197 ng/μl) and the percentage of RNA fragments greater than 200 bp was calculated (average 70.6%, all samples > 55%).

### NanoString PanCancer Immuno-Oncology (IO) 360

Extracted RNA was diluted to 30 ng/ul and 5 ul (150 ng) was combined with hybridization buffer and the Reporter CodeSet for the PanCancer IO 360 Panel (Nanostring) and incubated for 20 h at 65°C. The hybridized reaction was analyzed on an nCounter SPRINT Profiler (Nanostring). nSolver calculated normalization factors for each sample using raw gene counts and 14 housekeeping genes (Supplementary Table 1). Differential gene expression was calculated from normalized gene counts data and a false discovery rate with a Benjamini Hochberg multi-comparison test. The average counts for negative control probes was used for thresholding “positive” genes. After thresholding (<20 counts), a total of 666 genes passed filtration and were subsequently used for downstream analysis. The heatmap was generated using Clustergrammer [16], which is freely available at http://amp.pharm.mssm.edu/clustergrammer/. Raw gene counts are normalized using the logCPM method, filtered by selecting the genes with most variable expression, and transformed using the Z-score method.

nSolver advanced analysis tool was used for pathway analysis of the genes expressed in pre-versus post-NACT samples. The expression profile of the genes expressed was used to generate a pathway score for 25 different pathways (e.g., Hypoxia). The pathways scores were grouped and compared based on pre-versus post-NACT.

Genetic signature analysis was performed by Nanostring and previously described in [17–21]. Gene expression was normalized based on 14 housekeeping genes (Supplementary Table 1) and the log2-transformed gene expression values are multiplied by pre-defined weighted coefficient [18] and the sum of these values within each geneset is defined as the signature score.

### Reverse Phase Protein Array (RPPA)

Fresh snap-frozen tumor tissue was processed and analyzed by MD Anderson Functional Proteomics RPPA Core facility [22]. The RPPA (set167) included 449 unique antibodies, which were analyzed by Array-Pro Analyzer 6.3 then by SuperCurve 1.5.0 via SuperCurveGUI 2.1.1. The data were normalized based on the median. The data were centered based on the medians normalized to Log2 intensities.

### Transcriptional Factor Analysis

With the genes (n=18) that had a significant correlation to time to recurrence, we utilized PathwayNet Analysis to predict transcription factor relationships within the geneset [23]. Of this n=18 query genes, n=5 had no associated transcription factors; the resulting 13 query genes had 371 associated transcription factors (TFs) with any confidence. From this bipartite network, we derived a weighted projection onto the 13 query genes such that two nodes in the projection are connected by an edge weighted by the number of associated TFs they have in common. Under this projection, all TFs that associated with only one query gene are effectively removed, and higher weight edges can be interpreted as more reliable associations. Each node was then annotated by its “strength”, defined as the total weight of edges incident to it in the projection, i.e., the number of shared transcriptions factors with any other query gene. This operation produced a network with 13 nodes and 54 edges, accounting for 250 shared TFs.

### Tissue microarray (TMA)

A previously described [24] tissue microarray comprised of serous tumors from ovarian cancer patients treated at the University of Colorado (COMIRB #17-7788) was used to examine the tumor immune microenvironment via immunohistochemistry. The TMA consists of 109 primary HGSOC tumors collected pre-chemotherapy, 19 primary HGSOC collected post-chemotherapy, and 28 recurrent HGSOC tumors. Tumor regions selected by a board-certified pathologist, Dr. Miriam Post (Department of Pathology).

### Multi-spectral immunohistochemistry (IHC)

Multi-spectral IHC analyses were performed using Vectra Automated Quantitative Pathology Systems (Akoya Biosciences) as previously described [25]. The TMA slides were sequentially stained with antibodies specific for CD4 (T cells, 4B12, Leica Biosystems), CD8 (T cells, C8/144B, Agilent Technologies), FOXP3 (regulatory T cells, 236A/E7, Abcam), Granzyme B (effector T cells, GRB-7, Thermo Fisher Scientific), CD68 (macrophages, KP1, Agilent Technologies), and cytokeratin (tumor cells, AE1/AE3, Agilent Technologies). The entire TMA core was imaged using the 20x objective on the Vectra 3.0 microscope (Akoya Biosystems) and image analysis was performed using inForm software version 2.3 (Akoya), including tissue segmentation to define cells within the tumor regions (tumor-associated cells), cell segmentation to define cellular borders, and phenotyping to assign each cell to a phenotypic category. The TMA slides were also stained with IER3 antibody (Sigma Aldrich), imaged on the Vectra Polaris microscope (Akoya Biosystems), and analyzed with inForm software version 2.8 (Akoya) using a 4-bin algorithm to calculate histologic scores (H-scores).

### Statistical Analysis

Statistical analysis was performed using GraphPad Prism (Prism 8). Two-tailed student t-*test* or ANOVA were utilized to calculate a p-value. All quantitative data are graphed as mean with standard error mean (SEM). Hazard ratio calculated with Mantel Haenszel test. A calculated p-value of less than 0.05 was considered significant.

## RESULTS

Patients with advanced-stage HGSOC and extensive disease burden were treated with NACT, which provides a unique opportunity to investigate genetic and molecular change induced by carboplatin and paclitaxel (Figure 1A). Therefore, we examined the transcriptional profile of matched tumor samples from six patients pre- and post-NACT (total n=12). All six patients had extensive disease burden upon clinical presentation and were all treated with NACT with interval cytoreduction at The University of Colorado Cancer Center. These cases were selected for further analysis based on the availability of formalin-fixed paraffin-embedded (FFPE) blocks, snap-frozen tumor samples, and clinical information. The average time to disease recurrence was 243.5 days (range of 25 to 505 days); additional clinical attributes considered included genetic characteristics such as BRCA and homologous recombination (HR) status (Supplementary Figure 1A). Notably, based on CA125 levels at the time of diagnosis, there was not a clear relationship between disease burden and time to recurrence (Supplementary Table 2).

**Figure 1.**
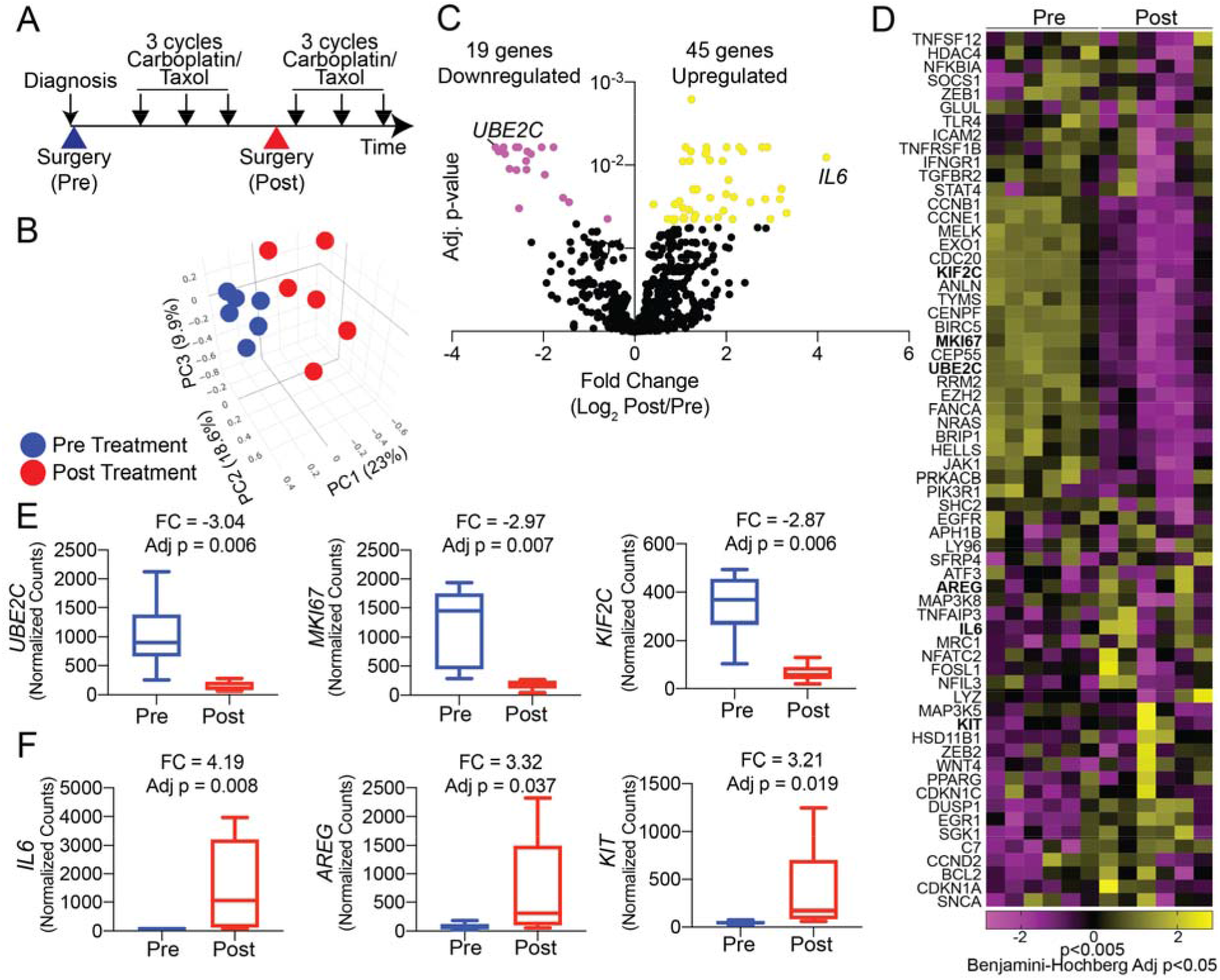
Chemotherapy-induced transcriptional dynamics. **A)** Graphic representation of NACT. Large arrowheads = surgeries, small black arrows = cycles of carboplatin/paclitaxel. **B)** Principal component (PC) analysis of the transcriptome of 770 genes in HGSOC tumors pre (blue) and post (red) chemotherapy. **C)** Volcano plot of differentially regulated genes. Yellow dots = Log2 FC<0 and adj. p-value <0.05. Pink dots = Log2 FC>0 and adj. p-value <0.05. **D)** Heatmap of 64 differentially expressed genes, generated with Clustergrammer from Nanostring data. **E)** Top 3 significantly downregulated genes following chemotherapy. FC = Log2 fold change Post/Pre. Adj. p-value calculated via Benjamini-Hochberg. Pre n=6 and Post n=6. **F)** Top 3 significantly upregulated genes following chemotherapy. FC= Log2 fold change Post/Pre. Adj. p-value calculated with Benjamini-Hochberg. Pre n=6 and Post n=6. For box plots the interior line indicates the mean and the error bars represent minimum and maximum values.

FFPE tumor tissue was subsequently identified and utilized for RNA isolation and transcriptomic analyses. We employed the NanoString platform to examine the expression of 770 highly annotated genes in the PanCancer IO 360 panel, which targets genes expressed by tumor cells and immune cells in the tumor microenvironment [21]. Linear regression of housekeeping genes between matched pre- and post-NACT tumors showed a significant correlation in 5 of 6 samples (Supplementary Figure 1B), confirming that housekeeping genes were comparable and could be used for normalization. While still significantly correlated, GTFB1064 samples demonstrated the highest degree of housekeeping gene variation between pre- and post-chemotherapy. Based on these successful quality-control benchmarks, we compared gene expression between matched samples, and across pre-versus post-NACT samples. A total of 666 target genes passed initial filtering (i.e. removal of genes with raw counts <20 for all samples) and were subsequently used for downstream analysis.

Principal component analysis of the 12 samples revealed a clear delineation between the pre- and post-NACT tumors (Figure 1B and Supplementary Figure 1C). This delineation suggests that there is a common or shared effect of chemotherapy on the tumor and tumor microenvironment (rather than chemotherapy causing primarily tumor-specific transcriptional changes). In total, 64 of 666 genes were differentially expressed between pre- versus post-NACT samples [adjusted (adj.) p < 0.05, Figure 1C–D and Supplementary Table 3]. The most strongly downregulated genes following NACT included *UBE2C*, *MIK67*, and *KIF2C* (Figure 1E and Supplementary Figure 1D). Downregulation of *MIK67* (Ki67, a proliferation marker) after NACT is consistent with chemotherapy targeting the dividing cells and causing cell cycle arrest. Conversely, the most strongly upregulated genes following chemotherapy were *IL6*, *AREG*, and *KIT* (Figure 1F and Supplementary Figure 1E). Chemotherapy has an established role in inducing IL-6 expression [7, 8], while upregulation of *AREG* (growth factor ligand Amphiregulin) and *KIT* (oncogenic receptor tyrosine kinase c-KIT) is consistent with activation of signaling through the epidermal growth factor receptor (EGFR) family of receptors. Pathway analysis of the gene expression profiles showed that, as expected, the Cell Proliferation pathway was significantly repressed in treated tumors (Table 1). Hypoxia response was the most significantly enriched pathway comparing pre-versus post-NACT gene expression, along with signaling downstream of AREG/EGFR and c-KIT (MAPK, PI3K/AKT, JAK/STAT) (Table 1). These data highlight that at the transcriptomic level, chemotherapy is targeting proliferating cells (as expected), but concomitantly activating oncogenic pathways. Intriguingly, elevated IL-6 and receptor tyrosine kinase (RTK) signaling by NACT may serve as a mechanism of resistance to chemotherapy.

**Table 1.** Pathway analysis of transcriptional profiles. Mean of pathway score, upper and lower limits of 95% confidence interval, p-value two-sided *t*-test, adj. p-value Benjamini-Hochberg multi-comparison correction.

| Pathway | Pre |  |  | Post |  |  | p-value | adj. p-value |
| --- | --- | --- | --- | --- | --- | --- | --- | --- |
|  | Mean | Upper Limit | Lower Limit | Mean | Upper Limit | Lower Limit |  |  |
| Hypoxia | -1.873 | -1.343 | -2.403 | 1.873 | 3.057 | 0.689 | 0.0000 | 0.0006 |
| PI3K-Akt | -2.764 | -1.875 | -3.653 | 2.764 | 4.678 | 0.850 | 0.0001 | 0.0006 |
| MAPK | -2.386 | -1.487 | -3.286 | 2.386 | 4.080 | 0.693 | 0.0001 | 0.0007 |
| Metabolic Stress | -2.721 | -1.662 | -3.780 | 2.721 | 4.743 | 0.699 | 0.0001 | 0.0007 |
| Cell Proliferation | 2.900 | 4.991 | 0.810 | -2.900 | -0.725 | -5.075 | 0.0006 | 0.0029 |
| Epigenetic Regulation | 1.159 | 1.984 | 0.335 | -1.159 | -0.188 | -2.131 | 0.0009 | 0.0036 |
| Apoptosis | 1.486 | 2.154 | 0.818 | -1.486 | 0.120 | -3.092 | 0.0013 | 0.0042 |
| Wnt Signaling | -1.634 | -0.444 | -2.825 | 1.634 | 3.100 | 0.169 | 0.0012 | 0.0042 |
| DNA Damage Repair | 1.543 | 2.803 | 0.282 | -1.543 | -0.187 | -2.898 | 0.0016 | 0.0044 |

To determine whether the transcriptional changes we observed post-NACT were associated with resistance to chemotherapy, we assessed whether the measured transcriptional changes between pre- and post-NACT samples correlated to time to recurrence. Of the 64 genes differentially expressed between pre- and post-NACT samples, 18 genes were correlated with time to recurrence (|r|>0.8) (Table 2). For instance, the magnitude of NACT-induced expression of the fibroblast growth factor receptor ligand *FGF9* correlated with time to recurrence (r= −0.990; i.e. high *FGF9* expression correlates with short time to recurrence). Signaling activated by FGF9 is consistent with the elevated PI3K/Akt and MAPK signaling observed in gene expression pathway analyses (Table 1). Since our transcriptional data were suggestive of a specific stress-like response to the post-NACT tumor microenvironment, we used the 18 genes correlated with time to recurrence to construct a transcriptional regulatory network of post-NACT gene expression, and hypothesized that this would identify stress response factors. In this network analysis, CEBPB (CCAAT enhancer binding protein beta; C/EBPβ) emerged as a central transcriptional hub driving a downstream network consisting of THBS1, IER3, OASL, IRF7, and CXCL5 (Figure 2A; Supplemental Table 4). This result implicates C/EBPβ as a central integrator of immune, inflammatory, and stress signaling in the EOC tumor microenvironment during NACT, which drives transcriptional programs mediating poor response to chemotherapy. Importantly, pathways activating C/EBPβ include growth factor signaling and IL-6 signaling [26], consistent with the genes and pathways identified above as upregulated in response to NACT.

**Table 2.**
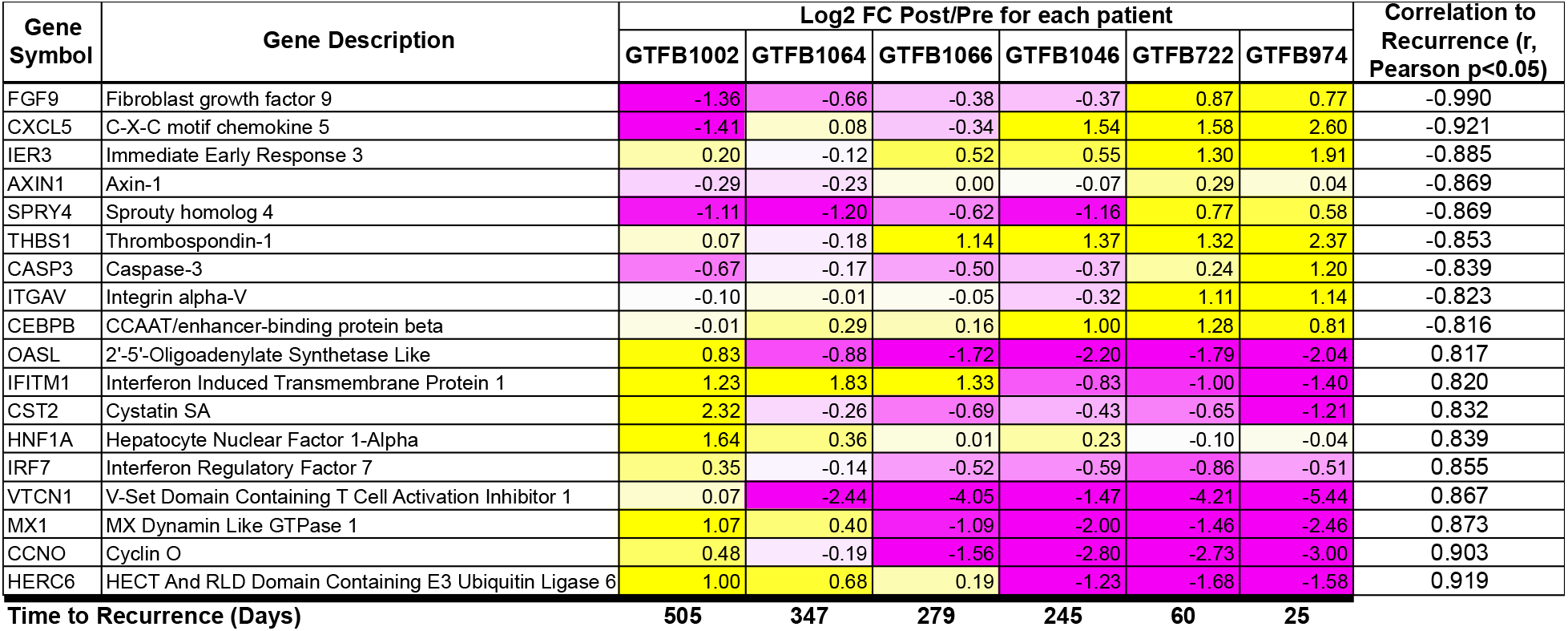
Gene expression changes that correlate to disease recurrence. Gene expression changes between pre- and post-chemotherapy Log2 fold change (FC) correlated with time to disease recurrence (last row) bold numbers. Yellow = upregulated, Magenta = downregulated. Pearson correlation analysis (|r|>0.8, p<0.05).

**Figure 2.**
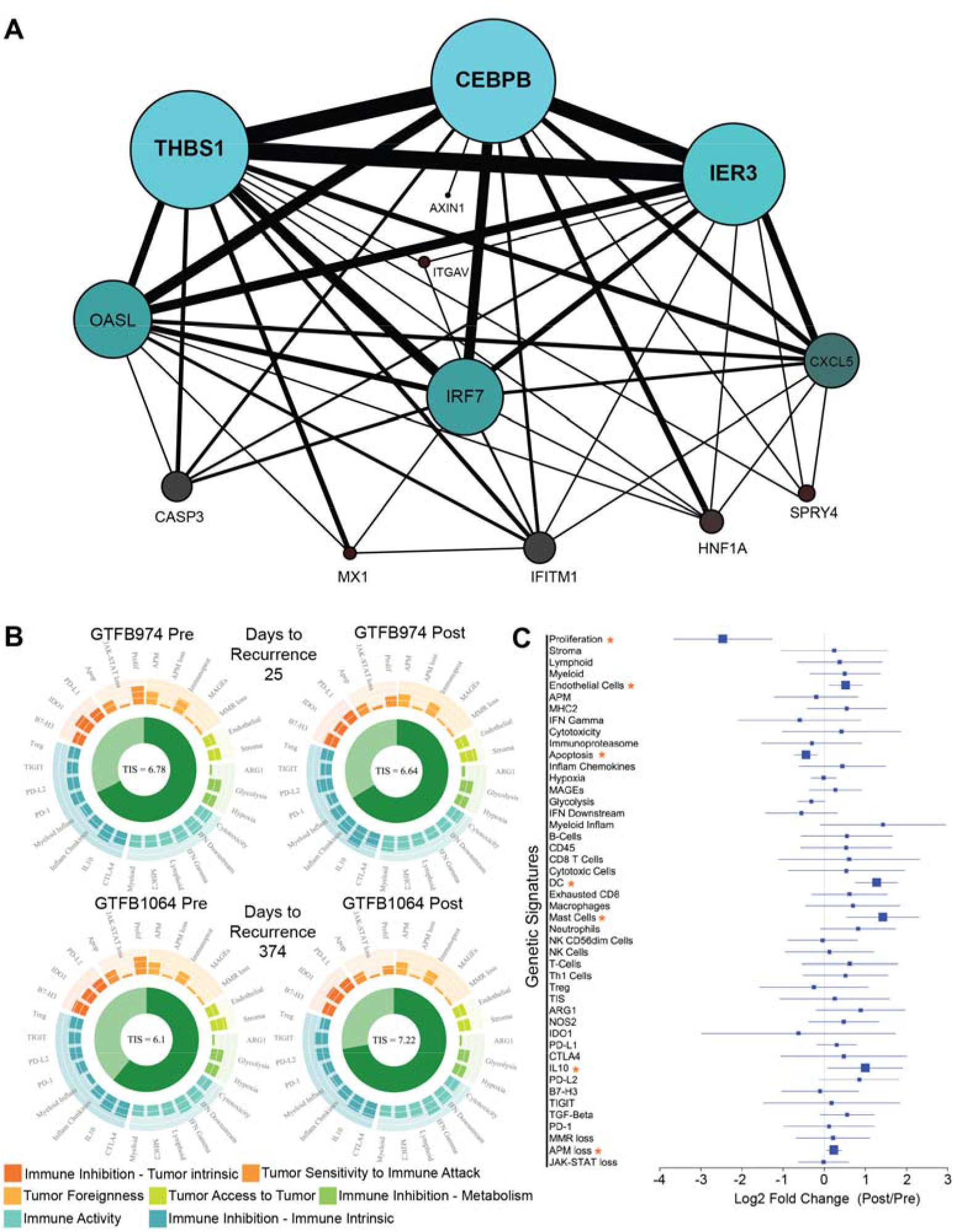
Transcriptional network and genetic signature analysis. **A)** Transcription factor relationship of 18 genes associated with disease recurrence (Table 2). Generated via PathwayNet data portal [23]. **B)** Utilizing the gene expression data genetic signature analyses was performed on all twelve samples. Shown are representative images of the genetic signature analysis from two sets of pre- and post-chemotherapy tumors (GTFB974 and GTFB1064). Each spoke on the circular plot indicates a unique genetic signature, such as proliferation, antigen-processing and presentation machinery loss, glycolysis. Genetic signatures are grouped by tumor immune microenvironment and characteristics of the tumor microenvironment. The Tumor Inflammation Score (TIS) is indicated at the center of the circular plot. **C)** Log2 fold change Forest plot of genetic signatures when grouped pre- and post-chemotherapy. Orange asterisks = p<0.05 and FDR<0.20.

We used a Nanostring panel in these studies with a series of pre-defined genetic signatures that extrapolate tumor microenvironment conditions, e.g. immune and stromal cell components, inflammatory signaling (e.g. tumor inflammation score), and immune activation state, based on gene expression (Figure 2B–C) [17–21]. Analyses using these genetic signatures, comparing pre-vs. post-NACT gene expression, identified a significant decrease (p<0.05) in “proliferation” and “apoptosis” signatures following chemotherapy. In contrast, signatures that were significantly upregulated (p<0.05) following chemotherapy include, antigen-processing and presentation machinery loss, IL-10, mast cells, dendritic cells, and endothelial cells (Supplementary Table 5). These data support our pathway analysis and suggest that increased inflammatory signaling driven by remodeling of the tumor immune infiltrate alter tumor cell signaling and ultimately drive chemotherapy resistance.

Since transcriptional analyses identified critical signaling pathways activated by chemotherapy, we directly assessed the activation of protein signaling cascades by RPPA. A limitation of this approach is the requirement for fresh frozen tissue samples, available from only two of the six patients treated with NACT (GTFB1064 and GTFB1066). In spite of this limitation, RPPA data closely aligned with Nanostring gene expression data for the 110 targets overlapping between the PanCancer IO 360 panel and M.D. Anderson’s standard RPPA platform (Figure 3A–C, Supplementary Table 6). Examining the fold change between the 110 overlapping targets, there was a significant positive correlation in both GTFB1064 and 1066 of the mRNA and protein fold expression changes (Post/Pre, Figure 3B–C). Consistent with suppression of proliferation by chemotherapy, transcriptional downregulation of *CCNB1* (Log2 fold change = −2.32, Adj. p=0.003) was paralleled by decreased Cyclin B1 levels in RPPA (Log2 fold change = −2.43) (Figure 3D–E). Consistent with signaling upregulation, *SGK1* mRNA and SGK1 protein were significantly upregulated post-NACT (mRNA Log2 fold change = 1.85, Adj. p=0.017; RPPA Log2 fold change = 0.76)(Figure 3F–G). Similarly, upregulation of *IL6* mRNA was paralleled by increased IL-6 protein levels (mRNA Log2 fold change = 4.19, Adj. p=0.008; RPPA Log2 fold change = 0.43)(Figure 3H–I). RPPA analyses also identified increased levels of phosphorylation of key signaling proteins, further supporting the activation of IL-6 and RTK signaling identified in transcriptional analyses. Levels of phosphorylated STAT1 (pY702), c-ABL (pY412), EGFR (pY1173), HER3 (pY1289), and IGFR (pY1135/36) were elevated post-NACT (Figure 3J–K). Notably, chemotherapy did not activate all receptor tyrosine kinases; phosphorylated HER2 (pY1248) and c-Met (pY1234/35) were unchanged following NACT (Supplementary Figure 2). These RPPA data closely parallel our transcriptomic data, together identifying increased IL-6 and activation of specific receptor tyrosine kinases post-NACT. Importantly, the RPPA data directly identify signaling cascades activated post-NACT, in particular JAK/STAT, which is a key downstream target of IL-6.

**Figure 3.**
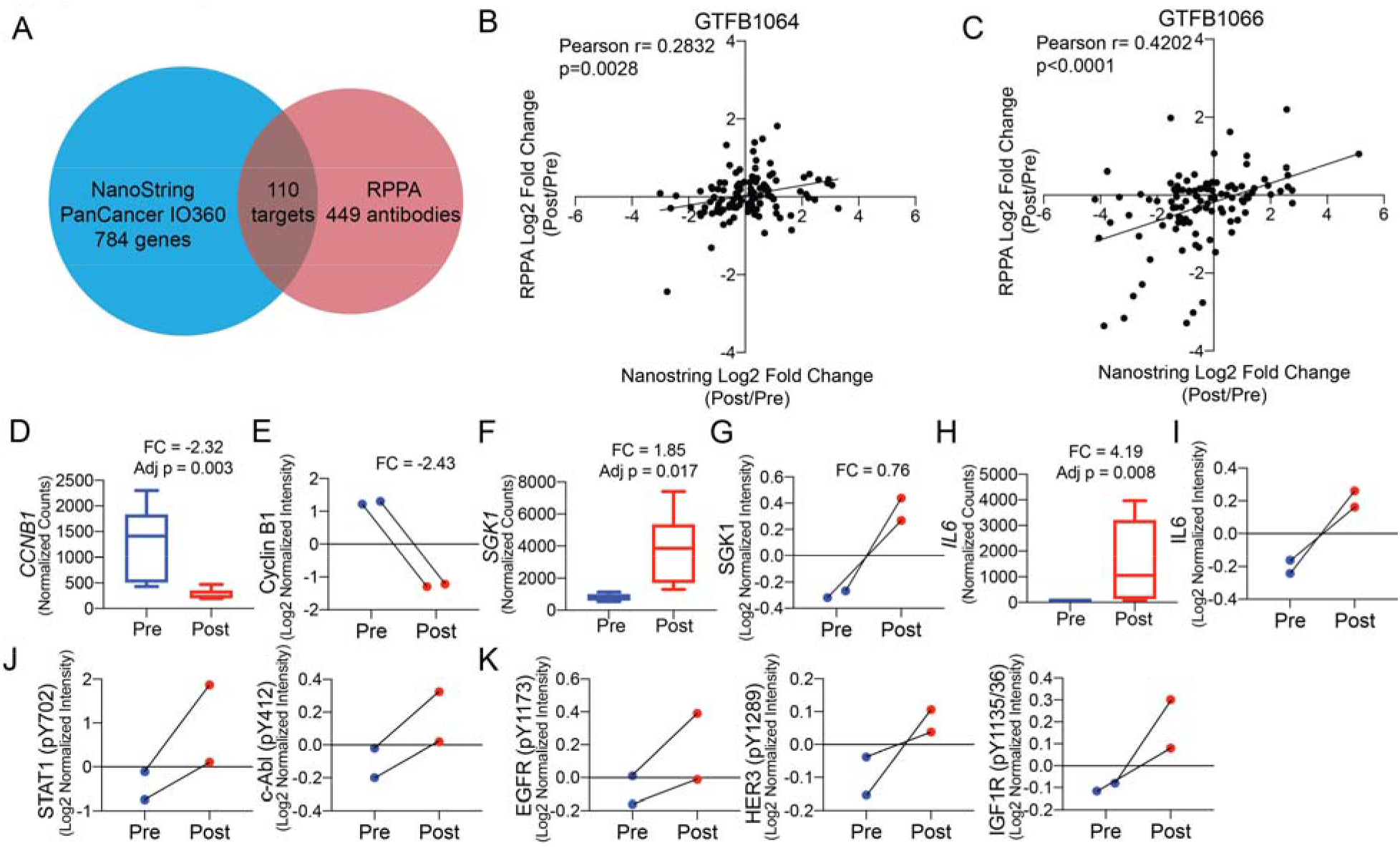
Reverse phase protein array of pre- and post-chemotherapy treated HGSOC tumors. **A)** Venn diagram indicating the overlap in targets from NanoString and RPPA. **B)** Scatter plot of log2 fold change (Post/Pre) of the overlapping targets of the RPPA (y-axis) and NanoString (x-axis) from GTFB1064. Pearson correlation calculated. **C)** Same as B, but with expression data from GTFB1066. **D)** Transcript counts of *CCNB1*. Pre n=6 and Post n=6. **E)** RPPA data of Cyclin B1 protein from pre- and post-treated tumors. **F)** Transcript counts of *SGK1*. Pre n=6 and Post n=6. **G)** RPPA data of SGK1 protein from pre and post-treated tumors. **H)** Transcript counts of *IL6*. **I)** RPPA data of IL-6 protein for pre and post-treated tumors. **J)** RPPA data of phosphoSTAT1 and phospho-c-Abl. Connecting lines indicate matched tumors. **K)** RPPA data of phosphorylated receptor tyrosine kinases. FC = Log2 fold change Post/Pre. Adj. p-value calculated via Benjamini-Hochberg. For box plots the interior line indicates the mean and the error bars represent minimum and maximum values. All other graphs error bars, SEM.

JAK/STAT signaling is initiated through the interaction of a cytokine receptor (e.g., IL6R) with its cognate ligand (e.g., IL-6), the latter of which may be secreted from the tumor or components of the tumor microenvironment, such as macrophages. As a surrogate for the tumor and associated microenvironment, we examined cytokine levels in ascites fluid from patients with HGSOC (most patients with advanced HGSOC present with ascites accumulation within the peritoneal cavity). Given that we found significantly increased IL-6 levels and JAK/STAT signaling activity post-NACT, we examined whether cytokine levels prior to chemotherapy predicted time to recurrence. Using primary ascites from 39 patients (pre-chemotherapy, ascites fluid is typically not available post-therapy), we used a multiplexed ELISA method to measure levels of the cytokines IFNγ, IL-10, IL-12p70, IL-13, IL-1β, IL-2, IL-4, IL-6, IL-8, and TNFαα Overall, IL-6, IL-8, IL-10, and IFNγ had the largest range of concentration (Figure 4A). We noted that IL-12p70 and IL-4 were significantly correlated (Figure 4B, Pearson r=0.9870, p<0.0001). Further, IL-6 positively correlated with five of ten cytokines. All of the cytokines were correlated to time until disease recurrence, which ranged from 1 to 66 months. Increasing concentrations of IL-6 or IL-10 in ascites correlated with a shorter time to disease recurrence (Figure 4C–D), suggesting that elevated cytokine signaling prior to or in response to chemotherapy may mediate resistance. Notably, there were several patients with low levels of IL-6 and IL-10 that recurred within ten months of the completion of primary chemotherapy (Figure 4C–D, red dots). These data demonstrate that while IL-6 (hazard ratio: 2.12) and IL-10 (hazard ratio: 1.52) concentrations predict time to disease recurrence, there is still a need to further elucidate the mechanisms underlying the tumor and immune microenvironment response to cytokine stimulation, especially in the context of chemotherapy.

**Figure 4.**
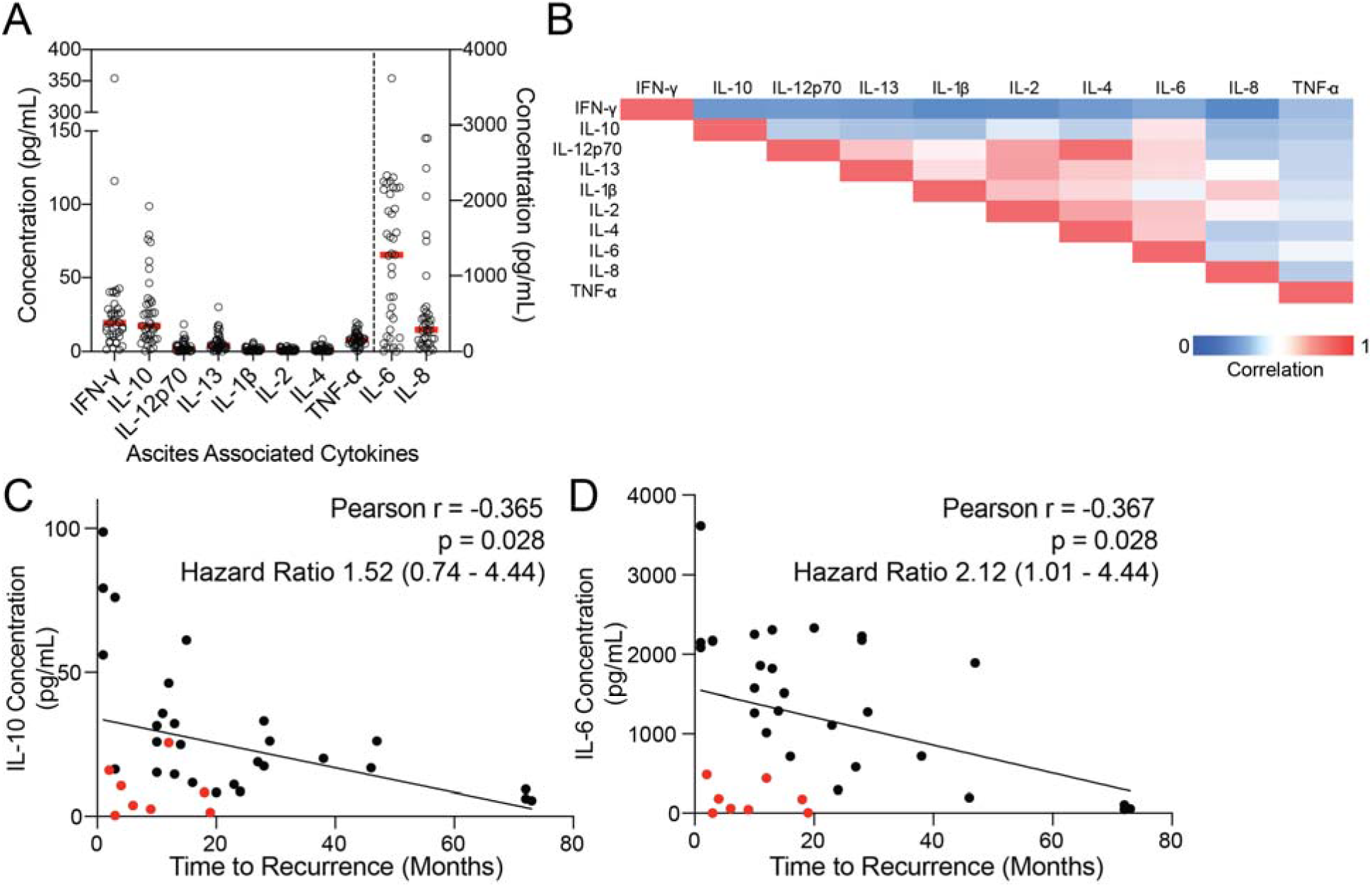
Cytokine profiling of primary patient-derived ascites. **A)** Ascites was collected from patients with HGSOC and was used for multi-plex ELISA from the indicated cytokines. Red bars = mean. Left y-axis include = IFN-y, IL-10, IL-12p70, IL-13, IL-1b, IL-2, IL-4, and TNF-a. The right y-axis includes IL-6 and IL-8. **B)** Correlation between the indicated cytokines. Red = 1 or strong correlation. Blue = 0 or weak correlation. (n = 39) **C)** Correlation between time to disease recurrence and IL-10 concentration. Hazard ratio calculated by stratifying data based on the median IL-10 concentration and using the Mantel-Haenszel test. **D)** Correlation between time to disease recurrence and IL-6 concentration. Red dots = short time to recurrence with low IL-6 concentrations. Hazard ratio calculated by stratifying data based on the median IL-6 concentration and using the Mantel-Haenszel test.

Transcriptomic and proteomic analysis of matched HGSOC samples pre- and post-NACT, coupled with cytokine profiling of pre-treatment HGSOC ascites, suggests elevated cytokine and/or RTK signaling prior to or in response to chemotherapy may drive resistance. In particular, our data implicate IL-6, consistent with previous studies. However, we noted that while IL-6 protein levels were associated with time to recurrence, a similar trend was not observed with *IL6* mRNA expression. Based on this, we hypothesized that IL-6 mRNA or protein levels alone may not sufficiently define activation of downstream cytokine signaling. Therefore, we compared the 18 genes that correlated with time to recurrence (Table 2) to *IL6* expression within the Nanostring dataset and the Cancer Genome Atlas (TCGA) ovarian cancer dataset (Supplementary Table 7). Between the two datasets, we observed that expression of *IER3* was the most positively correlated (Figure 5A–B) with *IL6*. Further, Figure 5C demonstrates that the magnitude of *IL6* induction following chemotherapy modestly correlated with time to disease recurrence. In contrast, the magnitude of *IER3* induction post-NACT strongly correlated with time to disease recurrence (Figure 5D), so we further examined IER3 protein expression via immunohistochemistry. Thus, using a tissue microarray (TMA) of 139 HGSOC tumors [24], which includes pre- and post-NACT primary tumors and recurrent tumors, we examined IER3 expression. When comparing pre- and post-NACT tumors and recurrent tumors, IER3 expression measured by histological score (H-score) was only significantly elevated in post-NACT tumors (p=0.0332; Figure 5E–F). Based on the median IER3 H-score, tumors were identified as having low (n=80) or high (n=39) IER3 expression and because tumor infiltration of T cells is a positive prognostic indicator [27, 28] we next used multi-spectral IHC to identify correlations of IER3 expression and immune cells in the tumor microenvironment. Specifically, we evaluated IER3 expression in relation to tumor-associated CD4+ T cells, CD8+ T cells, CD4+ FOXP3+ regulatory T cells (Tregs), CD68+ macrophages, pan-cytokeratin+ tumor cells, and Granzyme B+ T cells (Figure 5G). While tumor-associated macrophages and Tregs did not correlate with IER3 expression (Figure 5H–I), low IER3 expression significantly correlated with elevated tumor-associated CD4+ T cells (p=0.0087), CD8+ T cells (p=0.0502), and CD8/granzyme B+ T cells (p=0.0386) (Figure 5J–L). Taken together, these data suggest that regulation of IER3 and the composition of tumor immune microenvironment are interconnected.

**Figure 5.**
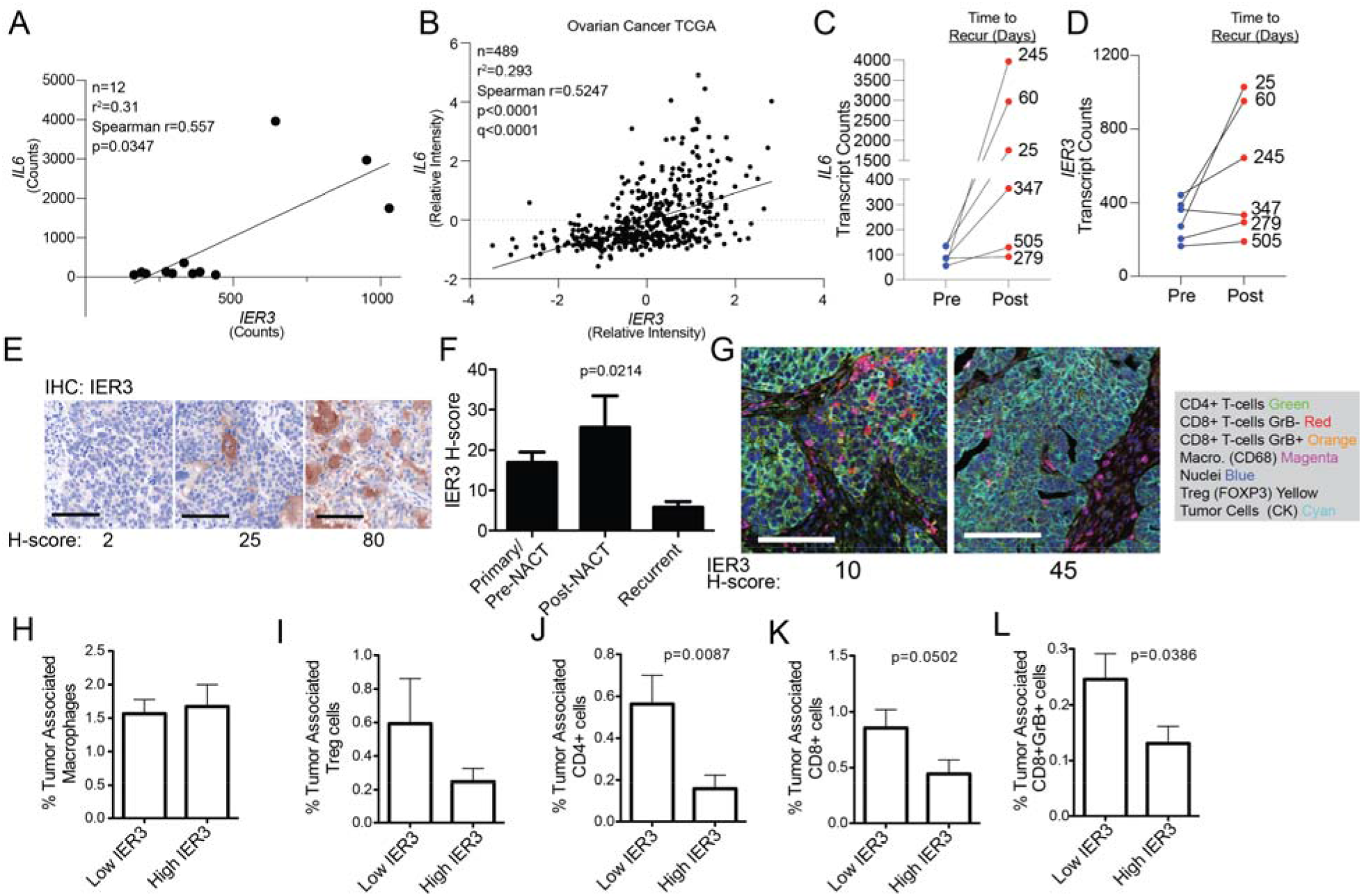
Transcriptional changes within matched tumors predict tumor recurrence. **A)** Correlation of *IL6* and *IER3* expression from pre and post-NACT samples. **B)***IL6* and *IER3* expression in 489 HGSOC tumors within TCGA. **C)** Change in *IL6* expression in pre- and post-treated tumors. Connecting lines indicate matching tumors. **D)** Change in *IER3* expression in pre- and post-treated tumors. Connecting lines indicate matching tumors. **E)** Representative images of IER3 IHC with associated histology score (H-score). Scale bars, 100 microns**. F)** IER3 histology score (H-score) for pre-(n= 85) and post-NACT (n=19) and recurrent (n=24) human HGSOC tumors. (p-value, one way ANOVA). **G)** Representative images of multi-spectral IHC of HGSOC tumors. Scale bars, 100 microns. Based on the median IER3 expression tumors were defined as “Low” (n=80) or “High” (n=39). The percentage of tumor-associated macrophages **(H)**, Tregs **(I)**, CD4+ T cells **(J)**, CD8+ T cells **(K)**, CD8/Granzyme B(Grb)+ T cells **(L)** was correlated to IER3 expression. (p-value, two-tailed t-test with Welch’s correction). Statistical test, unpaired *t*-test with Welch’s correction. Note: difference in n between panels G and H-L due to missing tissue cores. Error bars, SEM.

## DISCUSSION

A majority of patients with HGSOC will experience disease recurrence, with aggressive and chemo-resistant disease. Understanding how chemotherapy remodels the tumor and immune microenvironment is critical in predicting disease progression. To address this important issue in HGSOC biology, we utilized tumor tissue from patients that had received NACT, and evaluated treatment-induced molecular changes. We observed that chemotherapy attenuated cell cycle progression through the loss of cyclin B1 and Ki67, consistent with chemotherapy-induced cell cycle arrest. However, chemotherapy led to activation of several receptor tyrosine kinases (i.e., EGFR) and their downstream signaling pathways (i.e., JAK/STAT) associated with transcriptional programs of stress response and cell survival. Consistent with these data, we observed that NACT resulted in the upregulation of the inflammatory cytokine IL-6; elevated IL-6 and IL-10 levels at baseline also predicted disease recurrence. However, supporting the transcriptional and signaling network identified in our integrated analyses, expression levels of multiple CEBP/β target genes, in particular *IER3*, better predicts disease recurrence. Taken together, chemotherapy in HGSOC is paradoxically involved in promoting a more aggressive pro-tumor growth microenvironment, which remodels tumor cell signaling.

Chemotherapy results in significant tissue and cellular stress, which can lead to the activation of a multitude of effectors, including the transcription factor CEBP/β. The regulation of CEBP/β activity is highly complex, with multiple inhibitory and activating pathways (reviewed in [26]). Importantly, our analyses identify a positive feedback loop consistent with CEBP/β activation and upregulation of downstream target genes. As depicted in Figure 6A, although the response to surgery and chemotherapy is complex, other studies and our data suggest a convergence on CEBP/β transcription factor activity. CEBP/β has been independently identified as a prognostic factor for ovarian cancer and is linked to epigenetic regulation of multiple drug-resistance genes [29]. Based on CEBP/β activity in HGSOC, combination therapies that drive CEBP/β-dependence (e.g. NACT) and inhibit CEBP/β activation or transcriptional function (e.g. IL-6 blockade, RTK inhibition, or STAT3 inhibition) are potential approaches to improve the anti-tumor effect of chemotherapy [30–34]. However, our data suggest that in the context of surgery and chemotherapy, combinatorial therapies would likely be most beneficial in a maintenance setting following primary chemotherapy or NACT.

**Figure 6.**
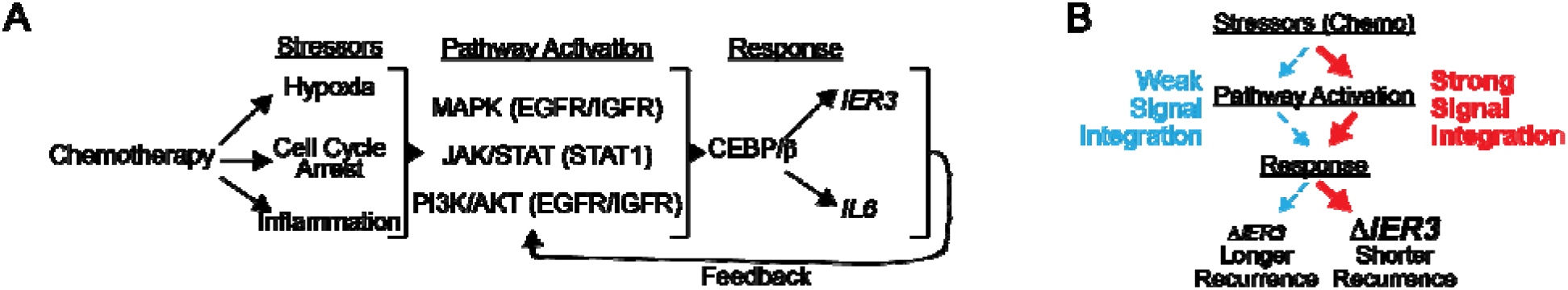
Working models of reported findings. Models for signal integration **(A)** and magnitude of signal intensity **(B)**.

By characterizing the tumor microenvironment over the course of treatment, unique information is obtained regarding the extent and magnitude of specific molecular changes. Previous studies that evaluated HGSOC tumors pre- and post-chemotherapy also noted that chemotherapy promoted an inflammatory response (i.e., increased IL-6), but the immunosuppressive environment prevented activation of anti-tumor immunity, suggesting a balance between inflammation and anti-tumor immunity. For instance, IL-6 and IL-10 cooperate to establish an immunosuppressive environment through expansion of myeloid-derived suppressor cells in ovarian cancer tumors [35]. Interestingly, in the ascites samples prior to chemotherapy, there was a subset of patients that had low IL-6 and IL-10 and a short disease-free interval, suggesting an immunologically “cold” tumor microenvironment and blunted anti-tumor responses to chemotherapy. Therefore, understanding the chemotherapy-induced effects that contribute to the maintenance of an immunosuppressive environment is critical. For instance, while IL-6 was upregulated in all of the treated tumors, IER3 was not, suggesting that tumors that upregulated IER3 are those competent to respond to IL-6. Consistent with differential IER3 inducton, the magnitude of chemotherapy-induced *IER3* upregulation was a superior predictor of disease recurrence compared to IL-6 alone. IER3 regulates extrinsic apoptosis and potentiates MAPK signaling. Further, in an inflammation-induced colon cancer mouse model, Ier3 knockout led to an exacerbated inflammatory environment, however the CEBP/β/IER3 signaling axis was not examined [36]. In our study, we observed significant correlations between IER3 expression and the tumor immune microenvironment, specifically that low IER3 correlated with increased CD4 and CD8 T cell infiltration. Since both *IL6* and *IER3* are key CEBP/β target genes (in addition to IL6 activating CEBP/β), the relationship between *IL6* and *IER3* suggests that increased levels of multiple genes activated by inflammatory/stress signaling post-NACT more accurately reflects pathway activation and therapy resistance. While IL6 likely plays a direct role in tumor progression as shown elsewhere, our integrated analyses suggest that tumor and immune microenvironments more poised to integrate and manage stress signals tend to be associated with a more aggressive, therapy resistant form of HGSOC (Figure 6A–B).

There are several limitations to our present study. The analysis only included six patients that qualified for NACT, which inherently suggests that their disease burden and stage were advanced. HGSOC frequently disseminates throughout the peritoneal cavity, which results in multiple sites of tumor colonization, and several studies have described different clonal patterns based on tumor sites such as omentum, ovary, or peritoneum [37]. The analysis is mostly one or two dimensional, and there is a significant need to increase the dimensionality of the datasets and to employ the emerging computational techniques to further define chemotherapy response further. Accounting for these limitations, our data and conclusions are still consistent and in close agreement with previous reports, and importantly, our study further expand on these previous reports by demonstrating how dynamic aspects of the tumor microenvironment can be exploited to improve patient outcomes in the future. Specifically we identified a novel, targetable IL-6/IER3 signaling axis that is enhanced following chemotherapy and promotes inflammatory signaling which may inform future lines of the therapy.

## ACKNOWLEDGEMENTS

Thank you to Dr. Philip Owens for his assistance with the Nanostring platform. We thank the Human Immune Monitoring Shared Resource of the University of Colorado Human Immunology and Immunotherapy Initiative for their expert assistance with the Nanostring and Vectra platforms. We acknowledge funding from The University of Colorado OB/GYN Academic Enrichment Fund (KB and TRK), Libations for Life (KB and TRK), and The University of Colorado Cancer Center Developmental Therapeutics Program (BGB and KB). This work was supported by grants from the NIH/NCI (BGB, R00CA194318-03; MJS, R00CA193734; JCC, U01CA231978), and Department of Defense Awards (JKR OC130212; BGB OC170228). KB is also supported by the Emily McClintock-Addlesperger Endowed Chair in Ovarian Cancer Research. The Functional Proteomics RPPA Core facility is supported by MD Anderson Cancer Center Support Grant # 5 P30 CA016672-40. Support of Core Facilities was provided by the University of Colorado Cancer Center Support Grant (P30CA046934).

## SUPPLEMENTARY INFORMATION

**Supplementary Figure 1.**
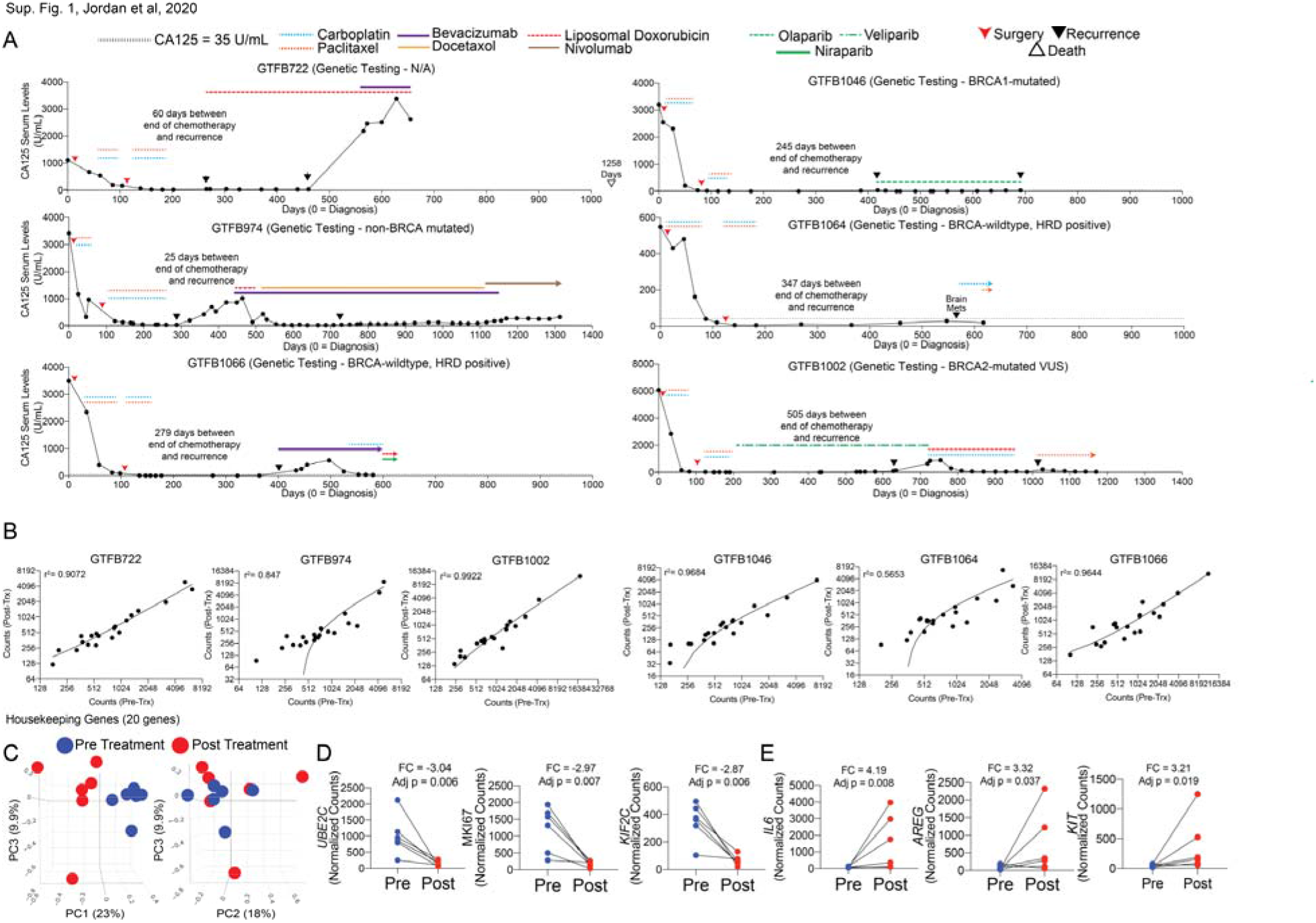
Disease progression for six patients with HGSOC and quality control for transcriptomics. **A)** Cancer antigen 125 (CA125, y-axis) tracks tumor progression over time (days, x-axis). **B)** Housekeeping genes of matched tumors for pre (x-axis) and post (y-axis) treated tumors. Linear regression of r = goodness of fit. Note: axes are log2 scale. **C)** Two additional angles of principal component (PC) analysis of the transcriptome of 770 genes in HGSOC tumors pre (blue) and post (red) chemotherapy. **D)** Three of the most upregulated genes in pre- and post-treated tumors. Connecting lines indicate matching tumors. **E)** Three of the most upregulated genes in pre- and post-treated tumors. Connecting lines indicate matching tumors.

**Supplementary Figure 2.**
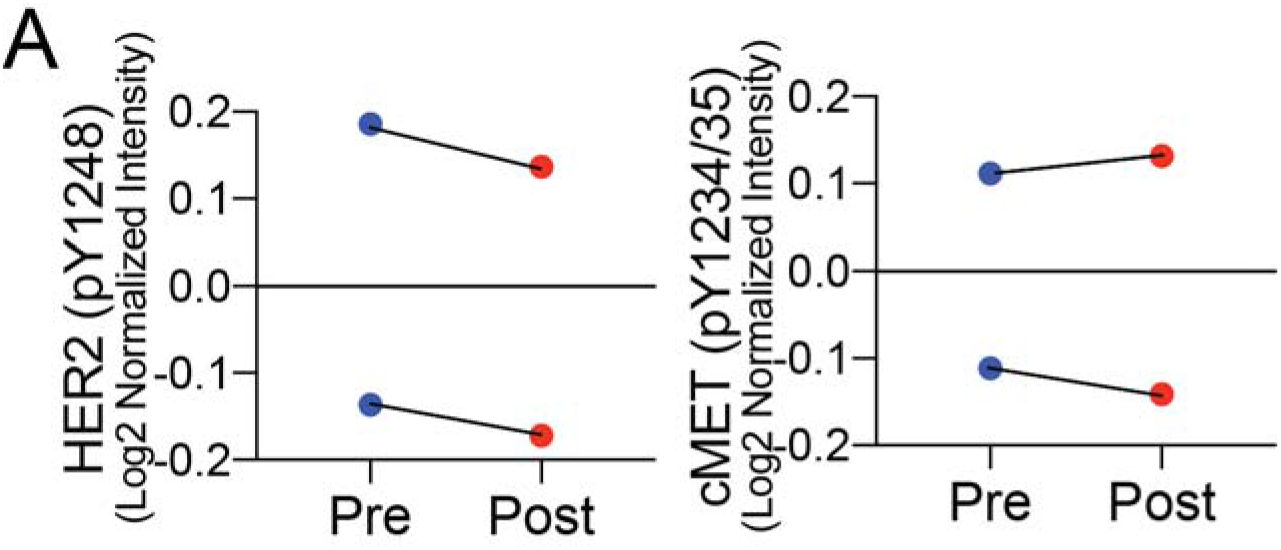
RPPA analysis of pre- and post-treated tumors. **A)** Log2 normalized intensities of receptor tyrosine kinases, HER2, and cMET. Lines indicate matching tumors.

### Supplementary Tables Legends

**Supplementary Table 1.** Normalization factors for each dataset calculated based on housekeeping genes.

**Supplementary Table 2.** Cancer antigen 125 (CA125) levels for each patient over time. Units = U/mL

**Supplementary Table 3.** Nanostring transcriptomic data for all genes include on the PanCancer IO 360 Panel.

**Supplementary Table 4.** Unthresholded transcription factor analysis from PathwayNet. Input genes are 18 genes associated with time to recurrence. Confidence values closer to 1 the higher the confidence.

**Supplementary Table 5.** Nanostring genetic signature analysis.

**Supplementary Table 6.** Reverse Protein Phase Array for GTFB 1064 and 1066 (pre and post-NACT).

**Supplementary Table 7.** Top 18 genes that correlated with *IL6* expression in Nanostring dataset (n=12) and TCGA dataset (n=489).

## Notes

The authors declare no potential conflicts of interest.

### Competing Interest Statement

The authors have declared no competing interest.

### Summary of Updates

Aaron Clauset's middle initial was incorrect on the first submission.

